# FGF21 Normalizes Plasma Glucose in Mouse Models of Type 1 Diabetes and Insulin Receptor Dysfunction

**DOI:** 10.1101/2021.01.04.425295

**Authors:** John L Diener, Sarah Mowbray, Waan-Jeng Huang, David Yowe, Jian Xu, Shari Caplan, Abhay Misra, Ankur Kapur, Jeffrey Shapiro, Xiaoling Ke, Xiaoping Wu, Avirup Bose, Darrell Panza, Min Chen, Valerie Beaulieu, Jiaping Gao

## Abstract

Fibroblast growth factor 21 (FGF21) is a member of the fibroblast growth factor (FGF) family of proteins. The biological activity of FGF21 was first shown to induce insulin independent glucose uptake in adipocytes through the GLUT1 transporter. Subsequently, it was shown to have effects on the liver to increase fatty acid oxidation. FGF21 treatment provides beneficial metabolic effects in both animal models and patients with obesity, type 2 diabetes mellitus (T2D) and/or fatty liver disease. In this paper, we revisited the original finding and found that insulin independent glucose uptake in adipocytes is preserved in the presence of an insulin receptor antagonist. Using a 40 kDa PEGylated (PEG) and half-life extended form of FGF21 (FGF21-PEG), we extended these *in vitro* results to two different mouse models of diabetes. FGF21-PEG normalized plasma glucose in streptozotocin-treated mice, a model of type 1 diabetes (T1D), without restoring pancreatic β-cell function. FGF21-PEG also normalized plasma glucose levels and improved glucose tolerance in mice chronically treated with an insulin competitive insulin receptor antagonist, a model of autoimmune/Type-B insulin resistance. These data extend the pharmacological potential of FGF21 beyond the settings of T2D, fatty liver and obesity.

## 1 Introduction

Fibroblast growth factor 21 (FGF21) is a member of the fibroblast growth factor family that mediates diverse metabolic actions in rodents, non-human primates and humans. Along with FGF19 and FGF23, it is an endocrine FGF as it can be secreted into the circulation. *In vitro*, FGF21 stimulates glucose uptake in adipocytes via increased expression of glucose transporter 1 (GLUT1) in an insulin-independent manner (Kharitonenkov et al., 2005). *In vivo*, FGF21 is a critical regulator of metabolism, and its expression is induced under conditions of both starvation and obesity (Badman et al. 2007, Domouzoglou and Maratos-Flier 2011, Fisher et al. 2011, Fazeli et al. 2015). When administered pharmacologically to both obese type 2 diabetic (T2D) mice and monkeys, FGF21 induces numerous beneficial metabolic effects, including body weight reduction, improved insulin sensitivity, glucose lowering (without evidence of hypoglycemia), increased high density lipoprotein-cholesterol (HDL-C), and decreased low density lipoprotein-cholesterol (LDL-C), triglycerides (TG), and hepatic TG content (Kharitonenkov et al. 2005, Kharitonenkov et al. 2007, Coskun et al. 2008, Veniant et al. 2012, Talukdar et al. 2016).

In T2D patients with increased body mass index, four-week treatment with FGF21 protein analogs results in mild weight loss but dramatic improvements in lipid profiles (Gaich et al. 2013, Talukdar et al. 2016). However, in these clinical trials the effects of FGF21 analogs on insulin sensitivity and glucose homeostasis were less profound, calling into question the translatability of this aspect of FGF21 biology to humans. Possible explanations for this lack of translatability include heterogeneity of the T2D patient population used in the studies and short duration of the studies (~ 4 weeks), which preclude the measurement of hemoglobin A1c (HbA1c), a more quantitative assessment of glucose control but requiring longer treatment for measurable changes. Given the ambiguous results observed in patients with T2D, we sought to assess the pharmacological effects of FGF21 on blood glucose under metabolic conditions significantly different from those previously investigated.

In this paper, we expanded on the initial observations of insulin independent glucose uptake (Kharitonenkov et al. 2005) in human and mouse adipocytes by FGF21. We further characterized the insulin independent actions of FGF21 in pharmacology studies using mouse models of type 1 diabetes (T1D) and in the setting of insulin receptor blockade, to establish that the insulin independent effects of FGF21 can translate from cultured cells to living organisms.

## 2 Materials and Methods

### 2.1 Protein Reagents

Insulin was purchased (Sigma). Carrier free human FGF21 was purchased (R&D Systems). The insulin receptor antagonist peptide, S961 (Vikram and Jena 2010), was custom synthesized (Bachem). PEGylated FGF21 analog, “FGF21-PEG,” was expressed, purified and PEGylated by Sandoz (Kundle, Austria). Briefly, the FGF21-PEG variant (aa 33-209: Q56E, D74H, Q82E, R105K, K150R, R154C, R159Q, S195A) was cloned into the modified *E.coli* expression vector pET30a, to generate in-frame fusions to a hexa-histidine tag followed by the N^pro^-EDDIE tag at the N-terminus of the FGF21 protein, expressed and purified using previously described methods (Achmuller et al. 2007). Freshly reduced protein was desalted and immediately PEGylated with 40 kDa branched maleimido-PEG reagent (NOF catalog # GL2-400MA from the Sunbright series). The PEGylated protein was finally purified by anion exchange chromatography and formulated at 50 mg/ml in a solution of Tris buffer (pH 7.7), sucrose and polysorbate 20. FGF21-PEG solutions were stored at 4°C and diluted in PBS for further use.

### 2.2 *In vitro* cell based assays

#### 2.2.1 Insulin and FGF21 dependent [^3^H]-2-deoxyglucose uptake in human adipocytes

Insulin and FGF21 induced [^3^H]-2-deoxyglucose (^3^H-2-DOG) uptake by human adipocytes was measured according to methods previously described (Kharitonenkov et al. 2005). Differentiated primary human adipocytes plated in a 96-well collagen-coated plate (Zenbio) were serum starved by incubating in 50 μl per well of Krebs ringer phosphate (KRP) buffer for 2 hours prior to treatments with either insulin (1 hour) or FGF21 (24 hours) in low glucose media (Invitrogen) supplemented with 0.5% heat-inactivated fetal bovine serum (FBS, Invitrogen), 100 IU/ml of penicillin, 100 μg/ml of streptomycin (Gibco) and 2 mM L-Glutamine (Gibco) at 37°C. Control cells (not exposed to insulin or FGF21) were treated with 1 μl/well of Cytocholasin B (an inhibitor of glucose transport) (Sigma) at a final concentration of 5 μg/ml for 30 min at 37°C in order to estimate background glucose uptake. ^3^H-2-DOG (20.6 mCi/mmol; 1 mCi/ml) was diluted 1:20 in 5.1 mM non-radioactive 2-DOG and 1 μl (0.05 uCi) was added to each well for 1 hour at 37°C. The plates were then washed 3 times with 100 μl/well of KRP-0.1% BSA buffer and 50 μl of 1% SDS was added to each well and plate shaken at medium agitation for overnight at room temperature. The following day, 200 μl of scintillation fluid (Optiphase Supermix, Perkin Elmer) was added per well, and the plates were sealed and radioactivity was measured using a beta microplate reader (Micro Beta Trilux). Cytochalasin B counts were averaged and subtracted from the control and treated wells. Data were analyzed and plotted as the fold change in glucose uptake for treated cells over controls. To measure the effects of insulin receptor antagonism on insulin or FGF21 dependent glucose uptake, cells were treated with the insulin receptor antagonist peptide S961 for 1 hour at 60 pM to 1 μM prior to addition of either insulin or FGF21.

#### 2.2.2 FGF21, human FGFR1c and human β-klotho dependent ERK phosphorylation assay

HEK293 cells stably co-expressing human FGFR1c and human β-klotho were maintained in DMEM media containing 10% FBS, 6.4 μg/mL blasticidin, and 0.64 mg/mL Geneticin at 37°C in 5% CO_2_. For the Extracellular-signal Regulated Kinase (ERK) 1/2 phosphorylation (pERK) assay, cells were plated into 384-well poly-D-lysine-coated plates (Becton Dickinson) at a density of 2.5 × 10^4^ cells per well in DMEM containing 10% FBS, and then incubated overnight at 37°C in 5% CO_2_. Overnight culture medium was removed and cells were then serum starved in 40 μl of Freestyle 293 media (Invitrogen) per well for 4 hours at 37°C in 5% CO_2_. FGF21 or FGF21-PEG were diluted in Freestyle 293 media and 20 μL of various concentrations of FGF21 or FGF21-PEG were added to each well and incubated for 10 min at 37°C in 5% CO_2_. Plates were emptied to remove the FGF21 containing media and were then immediately washed with 40 μL of ice-cold PBS per well (without Ca^++^ and Mg^++^, Invitrogen). The pERK AlphaScreen kit (Perkin Elmer) was used for measuring pERK activity. Seventy-five μL of ice-cold lysis buffer was added to each well and the cells were lysed by placing the plates on a shaker for 30 min at room temperature (RT). Four μL of cell lysate was transferred to a 384 well ProxiPlate (Perkin Elmer). Immediately prior to use, the reaction mix was prepared as per manufacturer’s instructions. The anti-rabbit IgG acceptor beads (Perkin Elmer) was also prepared according to the kit instructions. Five μL of the diluted acceptor bead mixture was added to 4 μL of cell lysate. The plate was sealed and incubated in dark for 90 min at RT. In reduced light conditions, streptavidin donor beads (Perkin Elmer) were diluted 40-fold with the dilution buffer provided and 2 μL of diluted donor beads were added per well to the cell lysate/acceptor bead mixture. The plate was sealed, and incubated in dark for 90 min at RT. Plate was then read on an EnVision 2104 multi-label reader (Perkin Elmer) using standard AlphaScreen settings. Dose-response data were graphed as pERK activity fold over basal versus protein concentration to determine EC_50_ values using GraphPad Prism.

#### 2.2.3 [^3^H]-2-deoxyglucose uptake in mouse 3T3-L1 adipocytes

3T3-L1 fibroblasts were purchased (ATCC). The cells were grown to confluency and were maintained in DMEM with high glucose (Invitrogen) supplemented with 10% FBS and 1% penicillin-streptomycin for an additional 4 days. They were then differentiated in the above media supplemented with 11 μg/ml insulin, 115 μg/ml IBMX (Sigma) and 0.0975 μg/ml dexamethasone (Sigma) for 3 days, after which the differentiation media was replaced with complete DMEM. The cells were distributed in 96-well plates one day before experiments. The next day, cells were treated with recombinant wild type FGF21 or FGF21-PEG overnight. After overnight treatment, the adipocytes were serum starved in 50 μl per well KRH buffer (4.5% Na Cl; 5.75% KCl; 7.85% CaCl2; 19.1% MgSO4; 0.25M HEPES, pH 7.5; 0.5% BSA and 2mM sodium pyruvate) for 2 hours. To estimate background uptake, a set of control cells were treated with 0.5 μl (final concentration 5 μg/ml) cytochalasin B for 15 min. ^3^H-2-DOG (20.6 mci/mmol, 1 mci/ml) was diluted 1:20 in 5.1 mM cold 2-DOG and 1 μl diluted 2-DOG per well was added to the cells and incubated for 5 min. The cells were washed with 100 μl/well KRH buffer three times. Fifty μl/well 1%SDS was added to cells and the cells were shaken for at least 10 min. Two hundred μl/well scintillation fluid was added and the plates were shaken for at least 2 hours and read in a beta-microplate reader (Micro beta Trilux). The data were plotted after calculating the fold over basal (no treatment) 2-DOG uptake (n = 4).

### 2.3 Diabetic mouse models

#### 2.3.1 Streptozotocin treatment in mice to induce type 1 diabetes

Twenty-two-week old male C57BL mice (Jackson Lab, Bar Harbor, ME) were housed four per cage in a standard light cycle room (light on from 6:00 a.m. to 6:00 p.m.) and given access to rodent chow (PicoLab Mouse Diet 20#5058, 21.6% kcal from fat) and water ad libitum. Diabetes was induced by an intraperitoneal (i.p.) injection of streptozotocin (STZ) (Sigma, St. Louis, MO, dissolved in 0.1 M citrate buffer at pH 4.5 and sterilized before injection) for three consecutive days at 70 mg/kg/day. When the mice were fully diabetic (19 days from the first STZ injection), they received either phosphate buffered saline (PBS) vehicle or FGF21-PEG two times a week (1 mg or 3 mg/kg, subcutaneous injection) for 26 days. One group of mice not treated with STZ was used as normal control and given PBS injections during the study. Experiments were conducted under an approved Institutional Animal Care and Use Committee protocol. All procedures in this study were in compliance with the Animal Welfare Act Regulations 9 CFR Parts 1, 2 and 3, and other guidelines (Guide for the Care and Use of Laboratory Animals, 1995).

Body weight and food intake were measured throughout the study. Plasma glucose, insulin, triglyceride (TG), free fatty acid (FFA) and β-hydroxybutyrate (BHB) concentrations and blood hemoglobin A1c (HbA1c) levels were measured under fed conditions. A glucose and arginine tolerance test (with an i.p. injection of a mixture of glucose at 1 g/kg and arginine at 0.6 g/kg) was given on day 23 after an overnight fast.

Glucose was measured using a glucose meter (Embrace, Omnis Health, Natick, MA). Insulin was measured using an ultra-sensitive mouse insulin enzyme-linked immunosorbent assay (ELISA) kit (Crystal Chem, Inc., Chicago, IL, cat# 90080). TG and FFA were measured using a highly sensitive fluorescent assay based on the horseradish peroxidase catalyzed oxidation of Amplex^®^ Red by hydrogen peroxide to resorufin. BHB was measured using an assay kit (Pointe Scientific, Inc.). HbA1c was measured using A1c Now+ Test Kit (Henry Schein, Melville, NY). Body composition was measured by EchoMRI-100 (Echo Medical Systems). Respiratory exchange ratio (RER) and energy expenditure were measured using TSE Systems (Bad Homburg, Germany) from day 16 to day 21 after mice had been acclimated in the system for 24 hours.

#### 2.3.2 Insulin receptor inhibition in mice to induce hyperglycemia and hyperinsulinemia

Twelve-week old male C57BL mice (Taconic, Germantown, NY) were housed individually in a standard light cycle room (light on from 6:00 a.m. to 6:00 p.m.) and given access to rodent chow (PicoLab Rodent Diet 20#5053) and water ad libitum. After an acclimation period, mice were implanted on the back with ALZET 2002 osmotic pumps (DURECT Corporation), containing PBS or insulin receptor antagonist S961 in PBS, delivering 10 nmol/week. Prior to the implantation, the pumps were loaded with PBS or S961 solution, soaked in sterile saline, and incubated at 37°C overnight. A dose of analgesics (Metacam at 5 mg/kg) was administered subcutaneously (s.c.) to the mice prior to anesthesia with isoflurane for pump implantation, and once daily for 2 days post the surgery. Three days after the pump implantation, mice implanted with S961 pumps were randomly assigned into two groups: S961 + PBS and S961 + FGF21-PEG, matched for their mean hyperglycemic levels. Mice in PBS and S961 + PBS groups were treated twice per week with s.c. injections of PBS. Mice in S961 + FGF21-PEG group were treated twice per week with s.c. injections of FGF21-PEG at 5 mg/kg. Body weight and food intake were measured during the study. Blood glucose and serum insulin concentrations were measured under fed and fasted conditions. An oral glucose tolerance test (OGTT) was performed on day 10 at 1 g/kg after an overnight fast. Blood glucose and serum insulin were measured before and during the glucose tolerance test at 0, 15, 30, 60 and 120 min of the OGTT. The study was terminated after the OGTT. The animals were euthanized with CO_2_. Cardiac puncture was performed to collect serum samples. Pancreatic tissue samples were collected, frozen in liquid nitrogen and stored at −80 °C for insulin content measurement. Glucose was measured using a glucose meter (Embrace, Omnis Health). Insulin was measured using an ultra-sensitive mouse insulin ELISA kit (Crystal Chem, Inc.). Frozen pancreatic samples were homogenized in buffer containing ice-cold acid/alcohol solution (75% ETOH/23.5% distilled water/1.5% 12N HCL). Protein content was measured using Pierce BCA protein assay kit (Thermo Fisher Scientific). Insulin content was measured using the insulin ELISA kit as mentioned above.

#### 2.3.3 Data analysis

Statistical analysis was performed using GraphPad Prism. Time course analysis was performed by a two-way analysis of variance (ANOVA) followed by a post-hoc test using Bonferroni’s method for each time point. Analysis among the groups without time course was conducted using a one-way ANOVA followed by Dunnett’s test. Data are presented as mean ± standard error of the mean (SEM). The level of statistical significance was set at p<0.05.

## 3 Results

### 3.1 FGF21 promotes glucose uptake in human adipocytes in the absence of insulin receptor signaling

We validated glucose uptake by human adipocytes using insulin to confirm cell function as expected (Figure 1a). We also replicated previously published results demonstrating FGF21 dependent glucose uptake by cells in the absence of insulin (Kharitonenkov et al. 2005) (Figure 1b).

**Figure 1.**
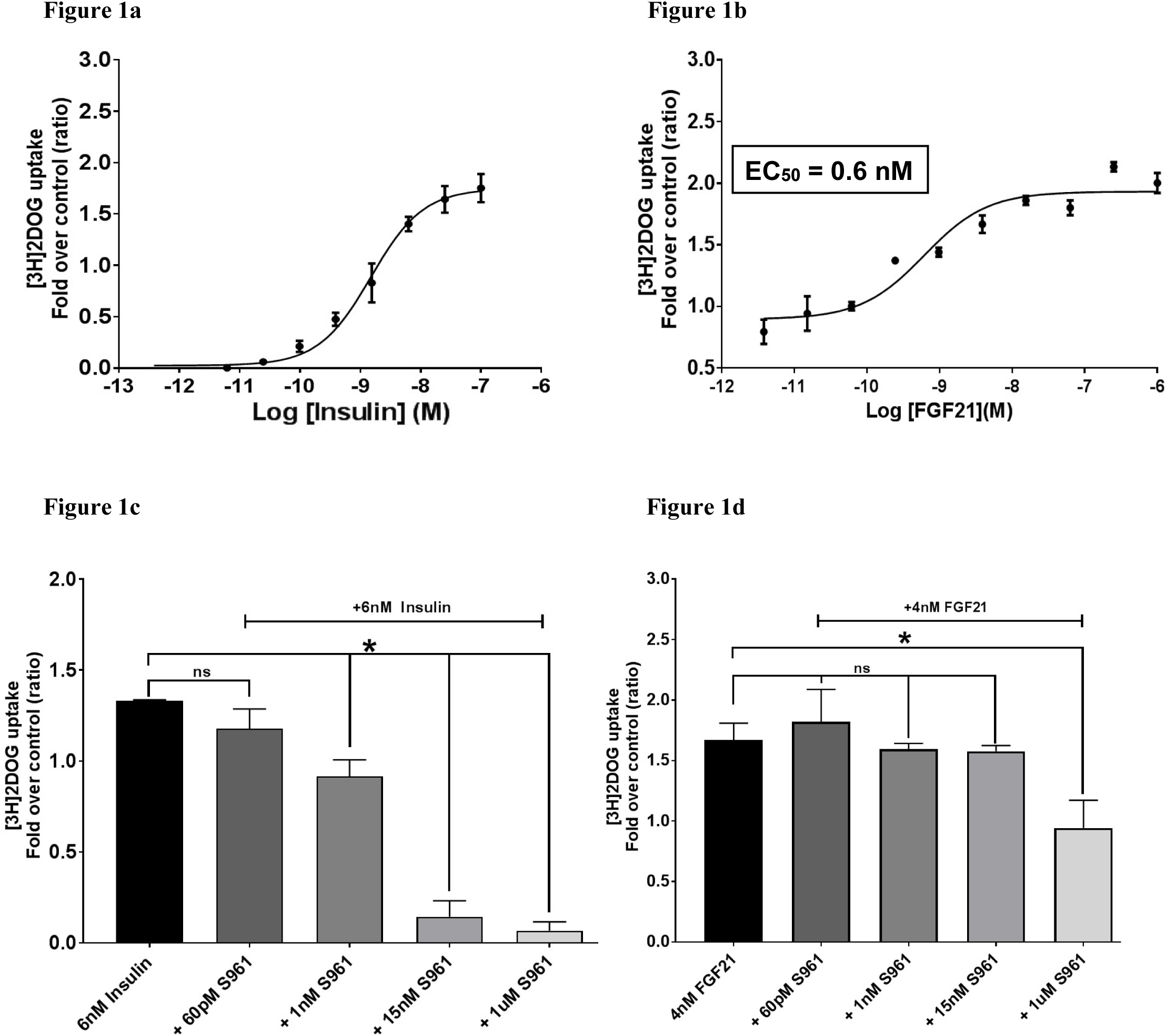
FGF21 increases glucose uptake in human adipocytes independent of insulin and with partial but not complete dependence on INSR. Glucose uptake by human adipocytes in the presence of insulin (Figure 1a) or FGF21 (Figure 1b). Glucose uptake by human adipocytes in the presence of low to high concentrations of S961with insulin (Figure 1c, S961 inhibits insulin-mediated 2-DOG activity at 1 nM (32%), 15 nM (90%), and 1 μM (>95%)) or FGF21 (Figure 1d, S961 inhibits FGF21-mediated 2-DOG activity only at 250 nM (44%, data not shown) and 1 μM (44% inhibition)). Asterisk (*) indicates significance (*p* <0.05) by one-way ANOVA.

To confirm if FGF21 promotes glucose uptake independently of insulin signaling, we assessed its activity in the presence of S961, a potent and selective peptide antagonist of the insulin receptor (INSR) that was derived from insulin (Vikram and Jena 2010). Pretreatment of human adipocytes with S961 at concentrations ≥ 15 nM for 1 hour at 37 °C completely abolished insulin-mediated glucose uptake in human adipocytes (Figure 1c). By contrast, treatment with 4 nM FGF21 induced glucose uptake in human adipocytes even in the presence of 1 µM S961, albeit at a slightly reduced rate (Figure 1d). These results indicate that FGF21 is sufficient to mediate glucose uptake in adipocytes under insulin receptor blockade and that, at most, FGF21 requires a small level of basal INSR activity for its full glucose uptake activity.

### 3.2 PEGylated FGF21 exhibits sufficient *in vitro* potency

To test whether FGF21 induces glucose uptake in the absence of insulin signaling *in vivo*, we engineered a PEGylated variant of FGF21 (FGF21-PEG), with an improved pharmacokinetic half-life and better exposure profile than unmodified wild type protein that enables twice weekly dosing. We compared *in vitro* activities of FGF21-PEG to that of FGF21 and confirmed the activity of FGF21-PEG by assessing phosphorylation of extracellular-signal regulated kinase (ERK1/2) in HEK293 cells expressing human FGFR1c and β-klotho. As shown in Figures 2a and 2b, FGF21 and FGF21-PEG activated pERK with EC_50_ values of 1.2 and 6.4 nM, respectively. In addition, FGF21 and FGF21-PEG induced glucose uptake in mouse 3T3-L1 cells with EC_50_ values of 0.5 and 3 nM, respectively (Figures 2c and 2d). These results indicate that the FGF21-PEG analog exhibits sufficient potency for *in vivo* testing.

**Figure 2.**
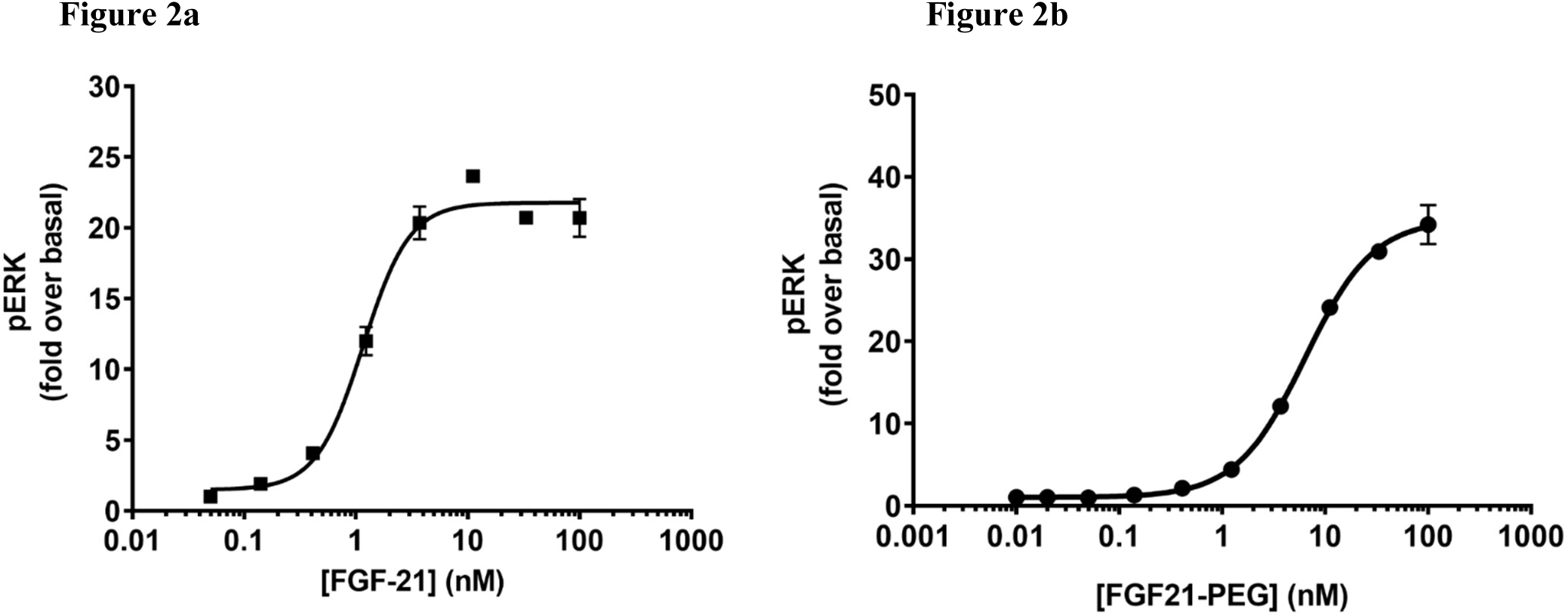
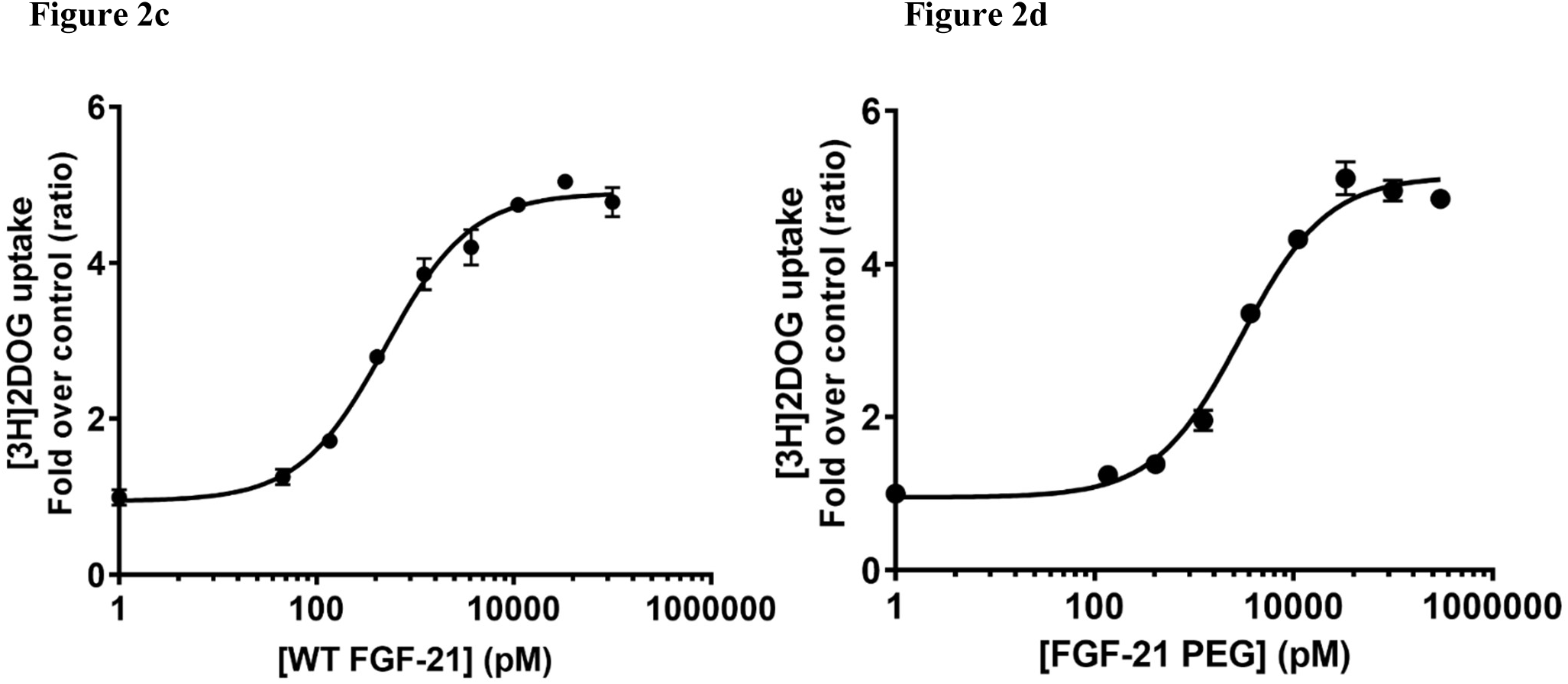
FGF21-PEG shows comparable potency to WT FGF21 on ERK phosphorylation in a human engineered cell line and glucose uptake in a mouse adipocyte cell line. ERK phosphorylation in HEK293 cells by WT FGF21 (Figure 2a) or FGF21-PEG (Figure 2b). Glucose uptake in mouse 3T3-L1 adipocytes by WT FGF21 (Figure 2c) or FGF21-PEG (Figure 2d).

### 3.3 FGF21-PEG normalizes glucose levels of streptozotocin-treated mice

Mice treated with the β-cell toxin streptozotocin (STZ) exhibit severe insulinopenia, hyperglycemia, dyslipidemia, ketosis and hyperphagia and are a commonly used rodent model of T1D. In adult C57BL mice treated with a high dose of STZ (70 mg/kg/day administered over the course of three consecutive days), β-cell regeneration is almost nonexistent and the animals remain insulinopenic. The effect of FGF21-PEG on T1D was evaluated under these conditions.

When severe diabetic hyperglycemia was induced 19 days following the initial STZ injection, the mice received either PBS vehicle or FGF21-PEG two times per week at 1 or 3 mg/kg by subcutaneous injections for 26 days. One group of mice not treated with STZ was used as a healthy normal control and given PBS injections during the study.

As shown in Figure 3a, profound hyperglycemia was observed in all STZ-treated groups prior to the treatment with FGF21-PEG or PBS vehicle. Treatment with FGF21-PEG resulted in a dose-dependent reduction of fed-state plasma glucose, with a significant decrease in plasma glucose from days 2 and 11 by 3 and 1 mg/kg, respectively. The 3 mg/kg treatment nearly normalized plasma glucose by the end of the study, compared to glucose levels of the control mice. In addition, FGF21-PEG treatment significantly reduced HbA1c (Figure 3b). In all STZ treated mice, HbA1c levels were significantly elevated prior to PBS vehicle or FGF21-PEG treatment. During treatment, HbA1c was further increased in the STZ-Vehicle group, while in animals treated with FGF21-PEG, HbA1c was either stabilized (1 mg/kg) or reversed (3 mg/kg), demonstrating a long-term improvement of glycemic control in a rodent model of T1D.

**Figure 3.**
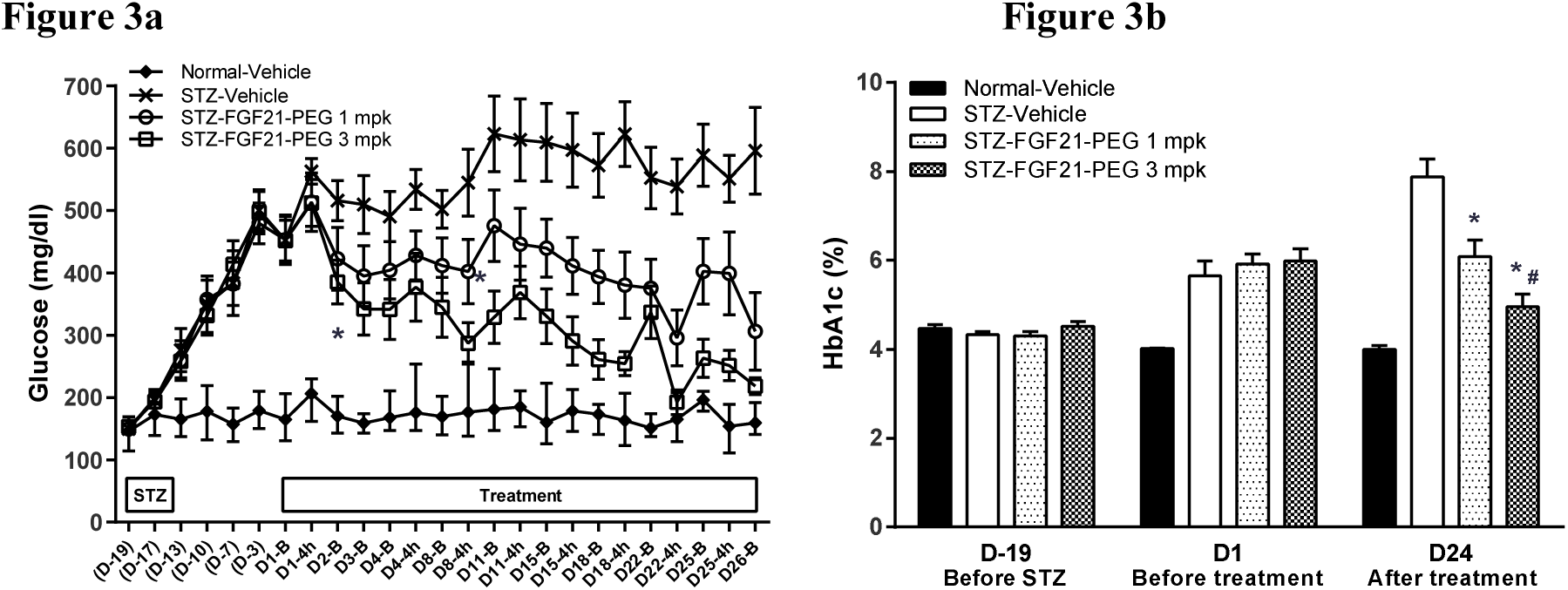
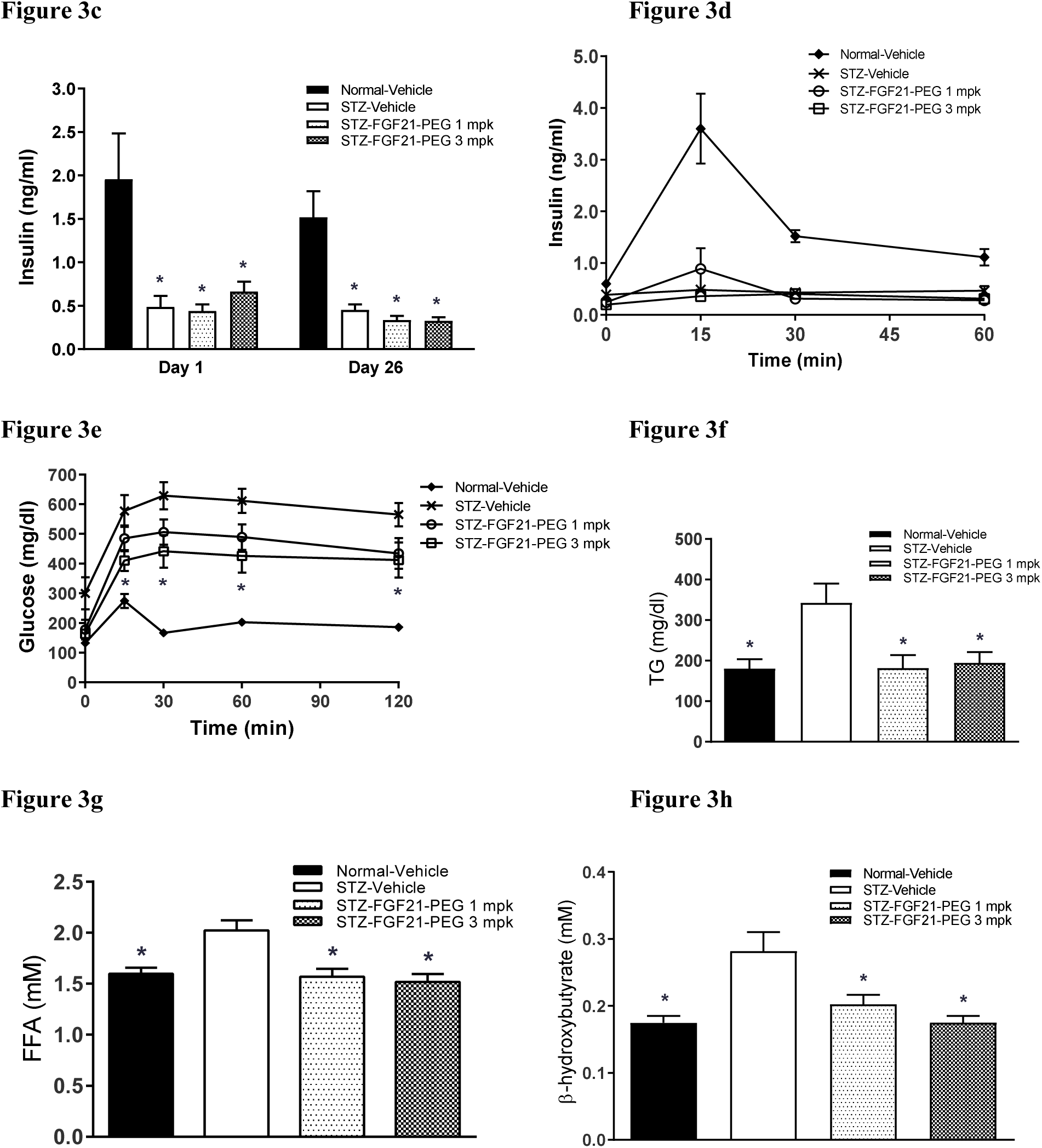
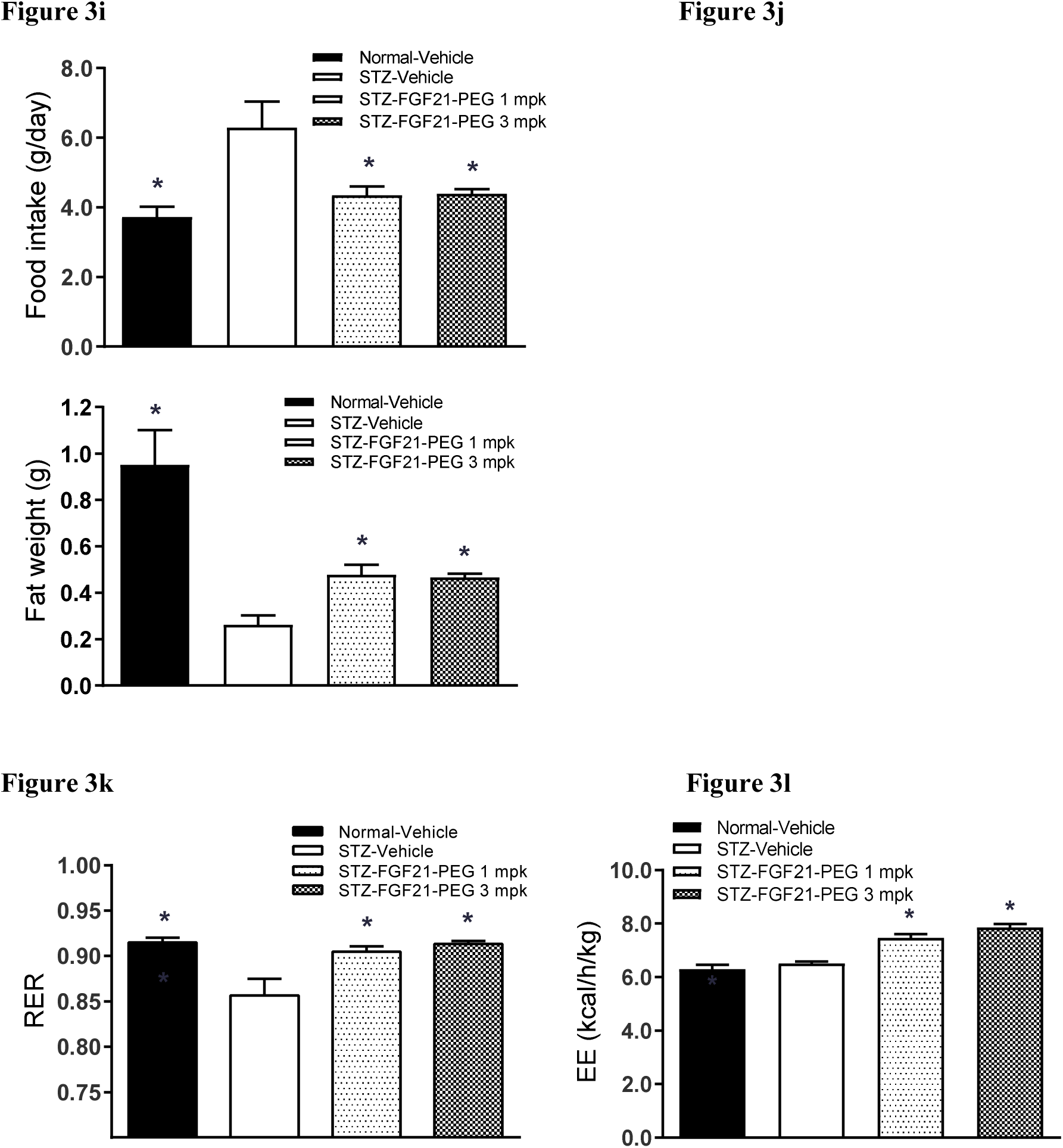
FGF21-PEG reverses hyperglycemia and HbA1c in a mouse model of T1D without restoration of insulin secretion. FGF21-PEG also normalizes metabolism on lipid and glucose utilization, shifting energy generation from fat to carbohydrate in T1D. Reduction of hyperglycemia (Figure 3a) and HbA1c (Figure 3b) by FGF21-PEG in STZ mice; plasma insulin levels at baseline (Figure 3c) and during a glucose and arginine tolerance test (Figure 3d); plasma glucose levels during a glucose and arginine tolerance test (Figure 3e); plasma levels of triglycerides (Figure 3f), fatty acids (Figure 3g), and β-hydroxybutyrate (Figure 3h); daily food intake (Figure 3i), epidydimal fat pad weight (Figure 3j), respiratory exchange ratio (RER, Figure 3k) and energy expenditure (EE, Figure 3l) in normal and STZ mice. Asterisk (*) indicates significance (*p* <0.05) by one-way or two-way ANOVA. #p<0.05 by one-way ANOVA, compared to STZ-FGF21-PEG 3 mpk on day 1.

FGF21 has been reported to have direct beneficial effects on pancreatic β–cells (Wente et al. 2006). To determine if the profound glucose lowering effect of FGF21-PEG in STZ mice was caused by restoration of pancreatic β–cells, plasma insulin levels were measured in STZ mice both prior to and after 26 days of treatment with PBS vehicle or FGF21-PEG. As shown in Figure 3c, plasma insulin levels were nearly depleted in all STZ treated groups and not restored after treatment with FGF21-PEG. Further, FGF21-PEG treatment had no effect on plasma insulin levels during a glucose and arginine tolerance test (Figure 3d). Thus, glucose lowering by FGF21-PEG in STZ-treated mice was not a consequence of the restoration of pancreatic β–cell function and insulin secretion, indicating that FGF21 mediates glycemic control independently of insulin signaling *in vivo*.

The glucose excursions of FGF21-PEG-treated animals during the glucose and arginine tolerance test were reduced significantly relative to vehicle-treated animals, but not completely normalized (Figure 3e). Thus, whereas FGF21-PEG exerts a profound effect on plasma glucose averaged over time, it is not sufficient to replace first-phase insulin secretion after an acute glucose spike.

Over the course of the study, the STZ-treated hyperglycemic mice became highly dependent on β-oxidation of fat stores for energy. As shown in Figures 3f-h, STZ mice treated with PBS vehicle exhibited elevated levels of plasma triglycerides, free fatty acids and β-hydroxybutyrate compared to normal mice. Treatment with FGF21-PEG normalized all of these biomarkers of fat mobilization and utilization, suggesting a shift from fat usage back to carbohydrate usage to generate ATP. We also assessed the effects of STZ and FGF21-PEG on whole body metabolism. Consistent with an inability to utilize carbohydrates for energy, STZ mice were hyperphagic and showed decreased epididymal fat mass and respiratory exchange ratio (RER) (Figure 3i-k). Treatment with FGF21-PEG decreased hyperphagia and increased RER, likely consequences of increased carbohydrate usage for energy. FGF21-PEG treatment led to a partial restoration of epididymal fat mass (Figure 3j). The increased energy expenditure with FGF21-PEG treatment (Figure 3l) together with the increased RER and partial restoration of fat mass indicate an improved carbohydrate utilization as a function of reversing T1D induced by STZ.

Taken together, these data suggest that FGF21-PEG could decrease plasma glucose in the absence of increased insulin secretion and promote the use of glucose as a major source of energy to improve glycemic control in individuals with T1D.

### 3.4 FGF21-PEG reverses hyperglycemia caused by insulin receptor blockade

Several rare insulin receptor disorders with either genetic (Donahue Syndrome, Rabson-Mendenhall Syndrome and Type-A Insulin Resistance) (Musso et al. 2004, Semple et al. 2011) or autoimmune (Type-B Insulin Resistance) (Malek et al. 2010) etiologies have been described. In all non-lethal cases, the severity of the disease is anti-correlated with residual insulin receptor signaling. In mice, homozygous whole-body knockout of the insulin receptor (INSR) is lethal within days of birth. However, several tissue-specific INSR knockout mouse strains have been generated and characterized, including liver (LIRKO), adipose (FIRKO) and pancreatic β-cells (BIRKO) (Kulkarni and Okada 2002). None of the tissue-specific INSR knockout mouse models fully replicate the effects of human loss-of-function mutations in INSR or autoantibodies to the INSR.

Given the profound glucose lowering effects of FGF21-PEG in insulin-deficient mice (Figure 3) and the preserved glucose uptake by FGF21 in adipocytes at the presence of insulin receptor blockade (Figure 1d), we then tested the effects of FGF21-PEG in mice with hyperglycemia induced by systemic insulin receptor blockade, which simulates insulin receptor disorders in people.

For this experiment we again used the insulin receptor antagonist S961 (Vikram and Jena 2010). Adult male C57BL mice were infused with PBS or S961 at 10 nmol/kg/week for 3 weeks. After one week of infusion, mice were treated with either 5 mg/kg FGF21-PEG or PBS vehicle by subcutaneous injections twice per week for the remaining two weeks. Mice that received S961 exhibited hyperglycemia within one week of the infusion (Figure 4a). Treatment with FGF21-PEG significantly reduced fed plasma glucose in S961-infused animals relative to vehicle-treated animals within 24 hours and normalized plasma glucose in less than two weeks (Figure 4a). Notably, when animals were overnight fasted on day 22, plasma glucose dropped to normal for both S961-treated groups (Figure 4d, time point = 0 min). Importantly, FGF21-PEG treatment following the overnight fast did not cause hypoglycemia by further decreasing plasma glucose levels.

**Figure 4.**
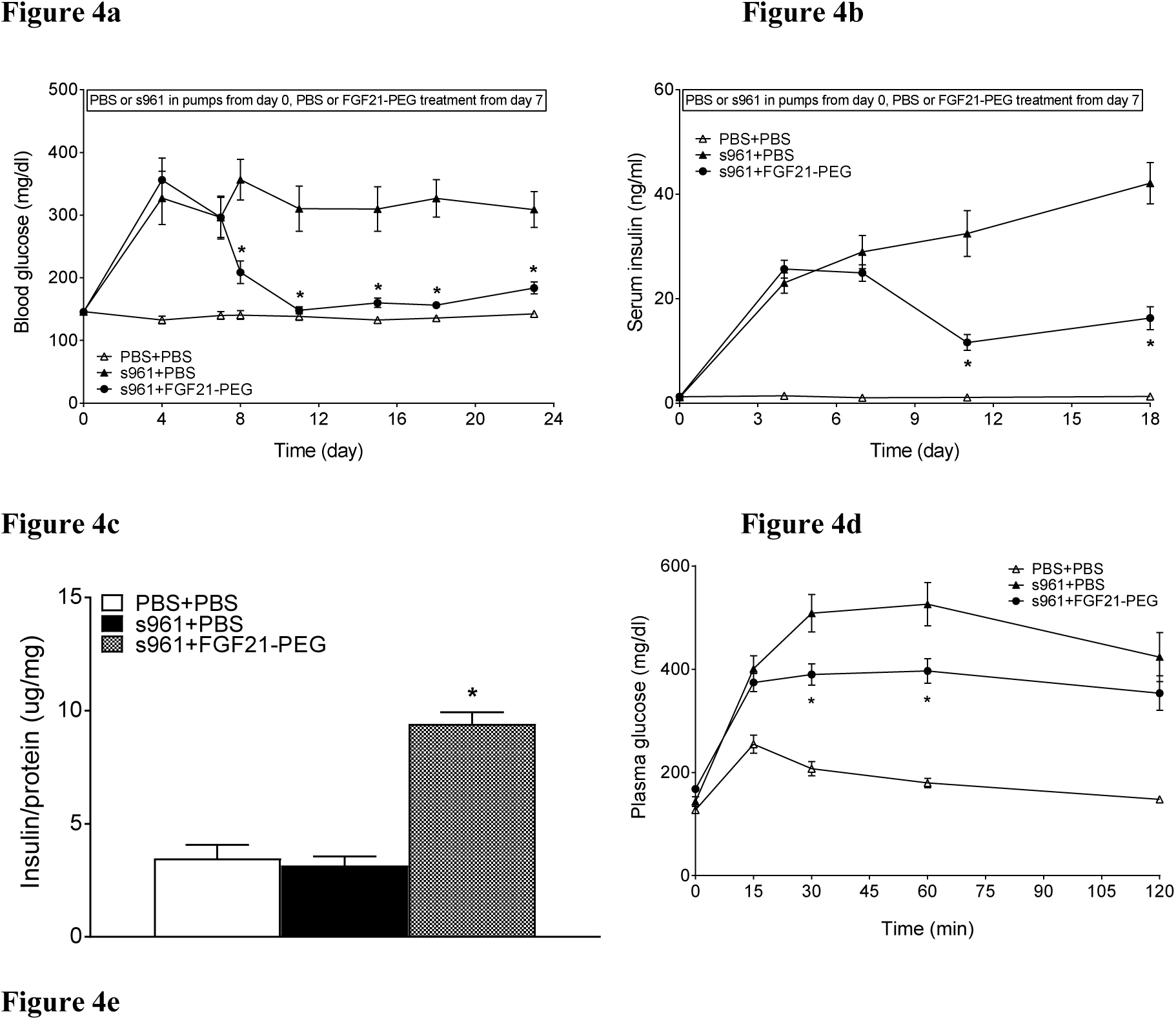
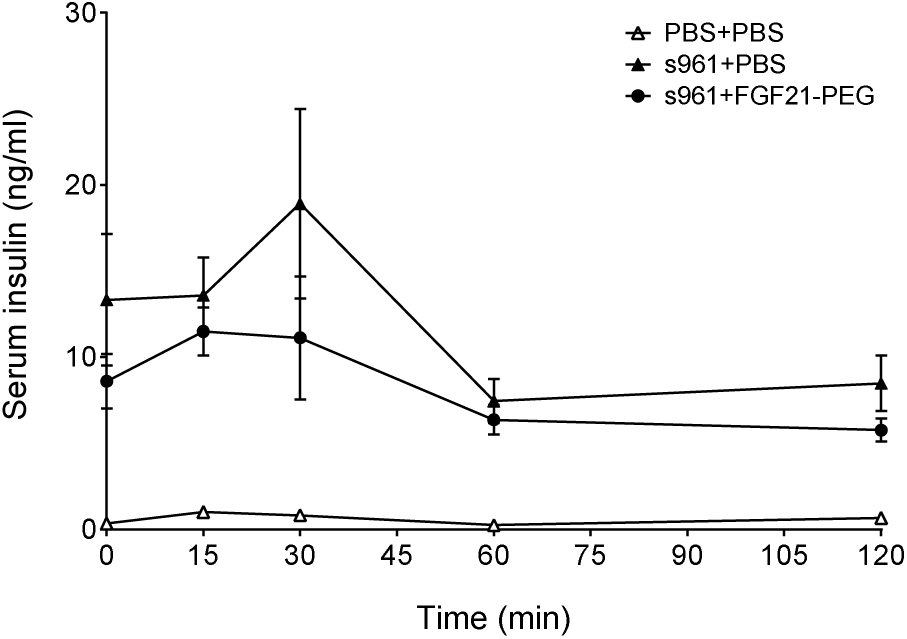
FGF21-PEG reverses hyperglycemia in a mouse model of insulin resistance and diabetes induced by S961. FGF21-PEG also preserves insulin content in pancreatic β-cells and improved glucose tolerance. Reduction of hyperglycemia (Figure 4a) and hyperinsulinemia (Figure 4b) by FGF21-PEG in S961-treated mice. Insulin content in pancreatic β-cells was increased by FGF21-PEG (Figure 4c), and glucose and insulin excursions were reduced by FGF2-PEG (Figures 4d and 4e). Asterisk (*) indicates significance (*p* <0.05) by one-way or two-way ANOVA.

As expected, all groups treated with insulin receptor antagonist S961 were hyperinsulinemic relative to the control group (Figure 4b). Treatment with FGF21-PEG mitigated the hyperinsulinemia in parallel with the reduction in hyperglycemia, and these animals had significantly higher pancreatic β-cell insulin content compared to vehicle-treated animals with or without blockade (Figure 4c). Taken together, these data suggest that an FGF21 therapy could have a significant β-cell sparing effect in patients with insulin receptor dysfunction. To test this hypothesis, a longer-term study would be required to demonstrate that chronic insulin receptor antagonism causes β-cell impairment.

FGF21-PEG reduced, but did not normalize completely, glucose and insulin excursions during an oral glucose tolerance test (OGTT) in S961-treated animals (Figures 4d and 4e). This result is consistent with our observations of STZ-treated mice (Figure 3e). Thus, FGF21-PEG can significantly reduce plasma glucose averaged over time in animals lacking insulin or insulin receptor signaling, but it is not sufficient to normalize glucose levels following an acute glucose spike.

## 4 Discussion

The potent metabolic pharmacological actions of FGF21 have been demonstrated in experiments using rodents, non-human primates, and, more recently, human patients. Transgenic mice with hepatic over-expression of FGF21 are lean, protected from high fat diet induced insulin resistance (Kharitonenkov et al. 2005, Inagaki et al. 2007) and demonstrate a substantial increase in longevity (Zhang et al. 2012). Furthermore, FGF21 protein administered to leptin deficient *ob/ob* mice, a rodent model of T2D, lowers blood glucose and triglycerides to near-normal levels without any evidence of hypoglycemia (Kharitonenkov et al. 2005). FGF21 also leads to decreases of body weight in this model (Coskun et al. 2008). Effects are also seen in diet-induced obese (DIO) mice, where exogenous FGF21 treatment leads to weight loss and increased energy expenditure potentially through the activation of brown adipose tissue and browning of white adipose tissue (Fisher et al. 2012). In diabetic monkeys, treatment with FGF21 or analogs also results in significant weight loss and improves lipoprotein profiles, including reduced LDL-C and increased HDL-C (Kharitonenkov et al. 2007, Adams et al. 2013). Likewise, treatment of patients with T2D with FGF21 resulted in modest weight loss and improvements in insulin sensitivity and lipid profiles (Gaich et al. 2013, Talukdar et al. 2016).

Despite the fact that the first reported activity of FGF21 was the insulin-independent induction of glucose uptake in adipocytes, subsequent characterizations of FGF21 have focused on its therapeutic potential for obesity and T2D. The studies described herein expand upon the original observation of insulin-independence, by assessing the utility of FGF21 in normalizing glucose levels in models of T1D with impaired or blocked insulin signaling.

We show that subcutaneous administration of a PEGylated FGF21 reverses hyperglycemia caused by STZ without any change in insulin secretion. Furthermore, we show that the FGF21-treated animals are able to use glucose for energy in addition to fat. These results are similar to those achieved using high doses of leptin in another model of T1D, the non-obese diabetic (NOD) mouse (Wang et al. 2010), and suggest that an FGF21 analog could benefit patients with established T1D, perhaps in conjunction with prandial insulin. Given that FGF21 does not induce hypoglycemia, an FGF21 therapy for T1D patients that significantly reduces insulin usage would confer significant safety benefits.

Our data also support evaluating the potential FGF21 analogs in patients with neutralizing antibodies to the insulin receptor, Type B insulin resistance (autoimmune insulin resistance), or other syndromes of insulin receptor deficiencies (*e.g.,* insulin receptor mutations such as Type A insulin resistance, Rabson Mendonhall Syndrome or Donohue Syndrome). The use of FGF21 was suggested as a possible treatment for these diseases in a manuscript describing the benefit of leptin therapy (~1.7% decrease in HbA1c) in 5 patients with Rabson-Mendenhall Syndrome (Brown et al. 2013). While a mechanism linking the metabolic effects of leptin and FGF21 has not been established, the shared benefits of both leptin and FGF21 in mouse models of T1D suggest that the benefits of leptin in patients with Rabson-Mendenhall Syndrome may also be achieved by treatment with FGF21.

## Acknowledgements

The authors would like to acknowledge Gerry Waters for helpful suggestions, and Allison Goldfine, Eleftheria Maratos-Flier and Daniel Denning for review of this manuscript.

